# Lateral ventricle volume trajectories predict response inhibition in older age - a longitudinal brain imaging and machine learning approach

**DOI:** 10.1101/468678

**Authors:** Astri J. Lundervold, Alexandra Vik, Arvid Lundervold

## Abstract

**Objective:** In a three-wave ~6 yrs longitudinal study we investigated if expansion of lateral ventricle (LV) volumes (regarded as a proxy for brain parenchyma loss) predicts performance on a test of response inhibition.

**Participants and Methods:** Anatomical trajectories of left (LH) and right (RH) lateral ventricle volumes across the three study-waves were quantified using the Longitudinal Stream in Freesurfer 5.3 and modelled using a linear mixed-effects (LME) algorithm. All participants (N = 74, mean age 60.7 yrs at inclusion, 48 females) performed the Color-Word Interference Test (CWIT). Response time on the third condition was used as a measure of response inhibition (RI) and divided into three classes (fast, medium and slow). The Extreme Gradient Boosting (XGBoost) algorithm was used for calculating the relative importance of selected LV volume features from the LME model in predicting RI class. Finally, the two most important extracted features were fed into a 10-fold cross-validation framework, estimating the accuracy, precision and recall of the RI class prediction.

**Results:** Four LME based features were selected to characterize LV volume trajectories: steepness of LV volume change and the LV volume at the time of inclusion, each from the right and left hemisphere. The XGBoost procedure selected the steepness measure from the right and the volume at inclusion from the left hemisphere as the two most important features to predict RI performance. The 10-fold cross validation procedure showed a recall, precision and accuracy score (.40 - .50) that were clearly above chance level.

**Conclusion:** Measures of the LV volume trajectories gave a fairly good prediction of response inhibition performance, confirming the role of LV volume as a biomarker of cognitive function in older adults. Future studies should investigate the value of the lateral ventricle volume trajectories as predictors of cognitive preservation or decline into older age.

## 1 Introduction

Normal aging is associated with morphometric changes in several brain regions and a corresponding decline in cognitive function, with trajectories of age-related changes that are characterized by individual differences [1]. Major events, biological and genetic factors through the lifespan obviously contribute to the heterogeneity observed in samples of older individuals [2,3,12], both regarding the rate of structural brain changes [5], the rate and extent of cognitive changes [6] and in brain-cognition relations [7,12]. In the severe end of the distribution, the most extensive parenchyma loss is associated with dementia, a syndrome defined by a severe decline in cognitive function [8]. On the other end of the scale we find so-called ”superagers” [9]. They show maintained cognitive function into old age [10], with a corresponding preservation of brain structure over time [11,12]. These findings support that age-related structural brain changes can act as strong predictors of cognitive abilities in old age, and emphasize the importance of taking individual differences into account in studies of associations between brain and cognition.

Several studies have linked changes in cognitive function to changes in specific regions of the brain (e.g., [13–16]). Prefrontal cortex is one such area, linked to both global aspects of cognitive function like fluent intelligence [17] and to specific measures defined within the concept of executive function (e.g., [18,19]). Executive function (EF) is of special interest in studies including older participants, as EF has been described as a hallmark of cognitive aging [20,21]. The concept of EF does, however, include several subcomponents. In the present study, we have focused on response inhibition (RI). RI is described as one of the core components of EF [22], and RI performance is known to be impaired as part of normal cognitive aging [23,24]. The close relation between response inhibition and fluent intelligence [25] and between fluid intelligence and various properties of brain structure [17] add to the interest of this EF subcomponent. The specificity of the relations between brain regions and RI and other subcomponents of EF is, however, still not clear. Inconsistent results are reported and can at least partly be explained by individual differences in age-related volume changes across different brain regions [26], but also by what Salthouse et al. [25] described as the ”ability impurity” of EF tests. In fact, subfunctions of EF are most likely dependent on multiple, interconnected brain regions [22]. In the present study, we will therefore not use volume changes in specific parenchymal brain regions as predictors of RI, but rather use trajectories of change in the lateral ventricle volumes as a more general proxy of age-related brain parenchyma loss.

The choice of LV volumes is further supported by studies describing the brain’s fluid-filled ventricles as a biomarker of the aging brain [27–29], and studies linking age-related ventricular expansion to changes in cognitive function at a subject-specific level [30–32]. A study by Todd et al. [33] showed a strong linear relationship between LV volume expansion and worsening of cognitive performance over a two-years period. The study assessed cognitive function by tests primarily designed to reveal symptoms of major neurocognitive disorders. Less is known about the longitudinal relationship between LV volume expansion and more specific measures of cognitive functions that are prone to normal age-related changes. The eight year longitudinal study by Long et al. [26] is an exception. The study assessed the co-evolution of volumetric brain changes and cognitive function in a large group of healthy older adults with tests defined as measures of different cognitive domains. The results showed volumetric reduction across several brain regions, and that faster cerebral atrophy and ventricular expansion were associated with rapid decline in performance on tests of verbal memory and executive function.

The study by Long and colleagues [26] motivated the present study to further investigate the ability of LV volume-derived biomarkers to predict cognitive performance. In the present study we restricted the evaluation of cognitive function to one test of RI: the third condition of the Color-Word interference (CWIT) test from the Delis and Kaplan Executive Function Scale (D-KEFS) [34]. We assume influence of processing speed and other fundamental abilities upon this test measure [24,35]. Moreover, its close relation to fluid intelligence [25] and various characteristics of the brain [17] make CWIT strongly susceptible to age-related changes. From this, we expect to confirm the expansion of ventricles that Long et al. [26] reported from their statistical mixed effects model, as well as an association between LV expansion and performance on CWIT. The main contribution of the present study is the extension of the statistical procedure used by most previous studies to include a machine learning framework, aiming at probabilistic prediction of RI performance from selected brain ”signatures” of LV volumes. More specifically, we first used a linear mixed-effect (LME) analysis similar to Long et al. [26] to select characteristics of the subject-specific trajectories of the left and right LV volumes. These characteristics were then included in a classification procedure, predicting individual belongings to one of three classes of performance level (fast, medium and slow) on the RI test. Feature importance was identified in the analyses using an extreme gradient boosting procedure. To obtain results that may generalize to other samples, we incorporated a small collection of both linear and non-linear classifiers in a *k*-fold cross validation scheme using multiple disjoint training and test sets drawn from our sample in order to estimate predication performance (i.e., accuracy, precision and recall). From previous studies we expected to find an age-related expansion of the LV volumes [36], a slower age-related expansion of LV volumes in women than in men [37–39], and that age would influence the response inhibition performance [23,24]. Our main hypothesis was that proper model based features characterizing the LV volume trajectories could act as good predictors of RI performance when included in a comprehensive classification framework.

## 2 Methods

### 2.1 Sample

The study included 74 healthy middle-aged and older subjects (48 females, 26 males; mean age 62.5 years at inclusion). They were all participants in a longitudinal investigation on cognitive aging, including neuropsychological and multi-modal brain MRI examinations. Subjects with a history of substance abuse, present neurological or psychiatric disorder, or other significant medical conditions when considered for inclusion, were excluded from participation. All subjects from the first study wave (*N* = 163) were invited to a follow-up study (wave 2) about three years later (*N* = 133), and to a third study (wave 3) about three years thereafter (*N* = 107).

Among the 107 participants of the third study-wave, 74 subjects provided successful and complete set of Freesurfer analyzed MRI data across the three waves and results from the cognitive test of response inhibition in wave 3. These subjects were included as the cohort of the present study.

An inspection of the neuropsychological test data from the three study-waves confirmed that none of the participants showed results indicating dementia. The test battery included two subtests from the Wechsler Abbreviated Scale of Intelligence (WASI, [40]) in the first study-wave to estimate intellectual function, and the Mini Mental Staus Examination (MMSE, [41]) in study-waves 2 and 3. All participants obtained a MMSE score equal to or above 25, and their mean IQ score was 117.1 (SD 10.2). None of the participants obtained a score on the second edition of the Beck Depression scale (BDI-II) [42] indicating a diagnosis of depression.

All participants signed an informed written consent form, and the Regional Committees for Medical and Health Research Ethics of Western Norway approved the three study waves.

### 2.2 Response inhibition

The total raw response-time (RT) score (in seconds) on the third condition of the CWIT [34] was included as the measure of response inhibition. In this condition, subjects are requested to name the colors of color-words printed in incongruent colors (e.g., the the word ”red” printed in green) as fast and correct as possible. From this, it is assumed that the participant has to inhibit the more automatic response to read the word, often referred to as the Stroop effect. In the two previous conditions of CWIT, the participants named a set of colours and read a set of color words. The third condition thus includes these two fundamental abilities, and their response time is also dependent on processing speed [35]. Trained research assistants administrated the test in a quiet room designed for a neuropsychological examination.

### 2.3 MRI acquisition and brain segmentation

Multi-modal MR imaging was performed on a 1.5 T GE Signa Echospeed scanner (MR laboratory, Haraldsplass Deaconess Hospital, Bergen) using a standard 8-channel head coil. Two consecutive T1-weighted 3D volumes were recorded from each subject (to improve SNR and brain segmentation) using a fast spoiled gradient echo (FSPGR) sequence (TE = 1.77 ms; TR = 9.12 ms; TI = 450 ms; FA = 7°; FoV = 240 × 240 mm^2^, image matrix = 256 × 256 × 124; voxel resolution = 0.94 × 0.94 × 1.40 mm^3^; TA = 6:38 min).

The same scanner (no upgrades) and T1-w 3D imaging protocol were used at each of the three study waves. Brain segmentation and morphometric analysis accross the three waves was conducted using the `FreeSurfer` image analysis suite, version 5.3 (documented and freely available online from https://surfer.nmr.mgh.harvard.edu). To extract reliable volume estimates and their trajectories (e.g. left and right lateral ventricles), the cross-sectionally processed images from the three study waves were subsequently run through the longitudinal stream [44] in FreeSurfer.

Specifically, an unbiased within-subject template space and image is created using robust, inverse consistent registration [43]. Several processing steps, such as skull stripping, Talairach transforms, atlas registration as well as spherical surface maps and parcellations are then initialized with common information from the within-subject template, significantly increasing reliability and statistical power [44]. As a consequence of the longitudinal processing stream and within-subject registration, the estimated total intracranial volume (eTIV) for a given subject remains fixed across the three study waves. Figs 1 illustrates the longitudinal MRI recordings (orig.mgz) and the corresponding FreeSurfer segmentations (aseg.mgz) from one of the participants at each of the three study waves. The age at the MRI examinations and corresponding left and right LV volumes are given along the time-line.

**Fig 1:**
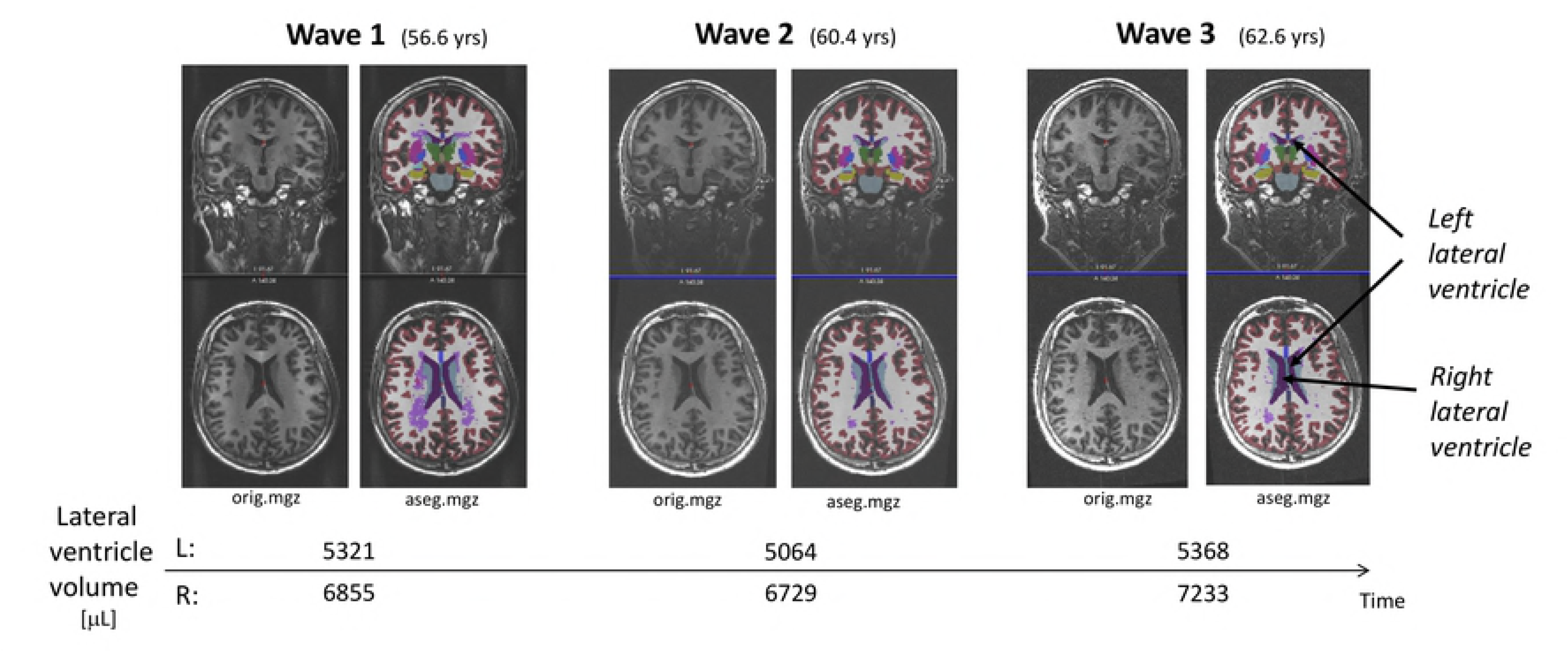
The longitudinal MRI recordings (orig.mgz) and the corresponding FreeSurfer segmentations (aseg.mgz) from one of the participants at each of the three study waves. The age at the MRI examinations and corresponding left and right lateral ventricle volumes are given along the time-line.

### 2.4 Statistical analyses

#### 2.4.1 Identification of individual trajectories of LV volume changes

Mixed effects modelling was used to identify measures of individual trajectories of LV volume change according to the following equation:

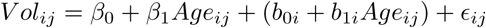

where *i* is subject (*i* = 1, …, *N* = 74) and *j* is wave (*j* = 1, …, *n* = 3). *Vol_ij_* - volume of left (right) lateral ventricle in subject *i* at wave *j* (response), and *Age_ij_* - age (in years) of subject *i* at wave *j* (predictor). *β*_0_ and *β*_1_ are fixed effects model parameters, *b*_0*i*_ and *b*_1*i*_ are random effects model parameters, and *ϵ_ij_*- is random residual errors, with zero mean and constant variance *δ* = *ϵ*2.

Two features were selected from the LME model to define the individual trajectories. The first is denoted *steepness of individual volume trajectory*, defined as the random slope effects *β*_1_. The second feature represents the LV volumes when included in the study, defined as the *deviance at wave 1 between subject*-*specific LV volume and the age*-*matched LV volume expected from the cohort fixed effect regression line*, Vdev (Figs 2). For each of these features, one is selected from the right and one from the left hemisphere. These four variables were included as predictors in further analyses.

**Fig. 2:**
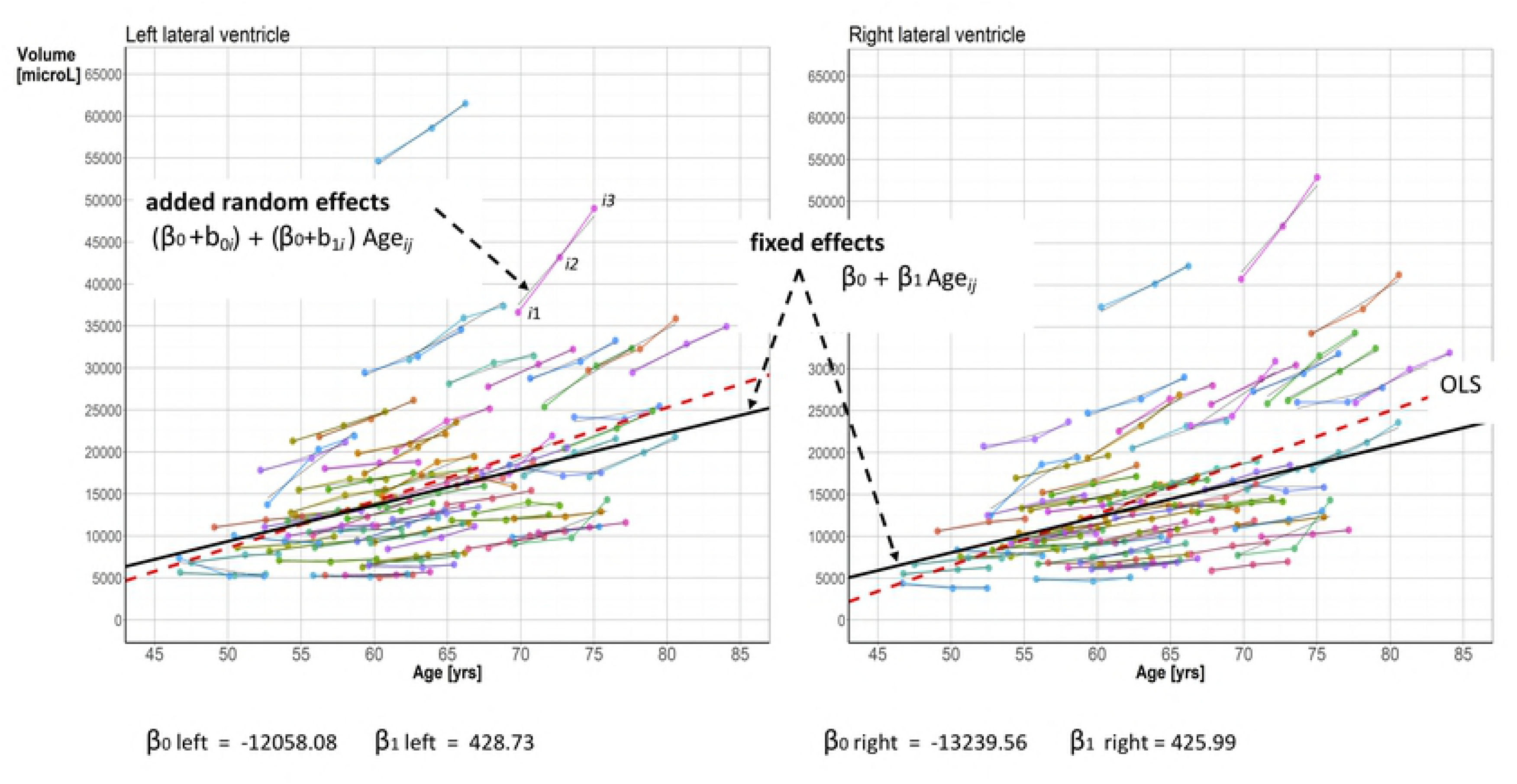
Illustration of he subject-specific measures of LV volume trajectories obtained from the LME analysis.

Results from descriptive statistics and correlation analyses were included to illustrate univariate associations between demographic data (age and gender), the selected measures of LV volume trajectories and response inhibition.

#### 2.4.2 Prediction of response inhibition

To study prediction of RI performance from the LV volume trajectories, we casted the problem as a classification task with balanced classes. To obtain such classes, we divided the participants into *fast* `(F)`, *medium* `(M)`, and *slow* `(S)` performers, using appropriate thresholds for the RI reaction time. By this we obtained equal prior probabilities of belonging to either F, M or S, (i.e, close to the same number of participants in each class). Technically we used the `qcut()` function in Pandas to compute the reaction time threshold values.

An extreme Gradient Boosting (XGBoost) procedure as provided by https://github.com/dmlc/xgboost was used to identify the order of importance of the measures of LV volume trajectories selected from the LME analysis (presented in the result part) in discriminating between the three levels of performance on the RI measure. This classifier implements a meta estimator that fits a number of randomized decision trees on various sub-samples of the dataset and use averaging to improve the predictive accuracy and control over-fitting.

To measure the quality of a split in a tree node, we used the `Gini` impurity criterion, and to aim at a statistically good results concerning the feature importance we specified `n_estimators` = 10000 as the number of trees in the forest. During the procedure, nodes are expanded until all leaves in the tree are pure.

#### 2.4.3 Prediction using *k*-fold cross validation framework

From the feature importance analysis, the top two ranked predictors selected to represent the trajectories of the LV volumes were included for a comprehensive classification study. We used *k*-fold cross validation to assess the prediction properties (accuracy, precision, recall). This was applied by including a linear (multinomial logistic regression = MLR) and a non-linear classifier (multi-layer perceptron = MLP, with three layers) and a voting classifier.

For the performance assessment on each fold, we used the *accuracy*_*score* (the ratio of correct classifications), *precision*_*score* (the ratio *tp/(tp+fp)*, where *tp* is the number of true positive and *fp* number false positives), *recall*_*score* (sensitivity, the ratio *tp/tp+fn)* where *fn* is the number of false negatives. All these measures were generated from `sklearn.metrics`.

These supervised data-driven machine learning analysis were implemented in Jupyter notebooks using Python (3.5.4), Numpy (1.12), Pandas (0.20), Statsmodels (0.8), XGBoost (0.6), Scikit-learn (0.19), rpy2 (2.8.5) and Matplotlib (2.0) for producing Figs 3 and 4. Our Jupyter notebook for computing feature importance and classification with *k*-fold cross-validation will be available on GitHub [address TBA].

**Fig. 3:**
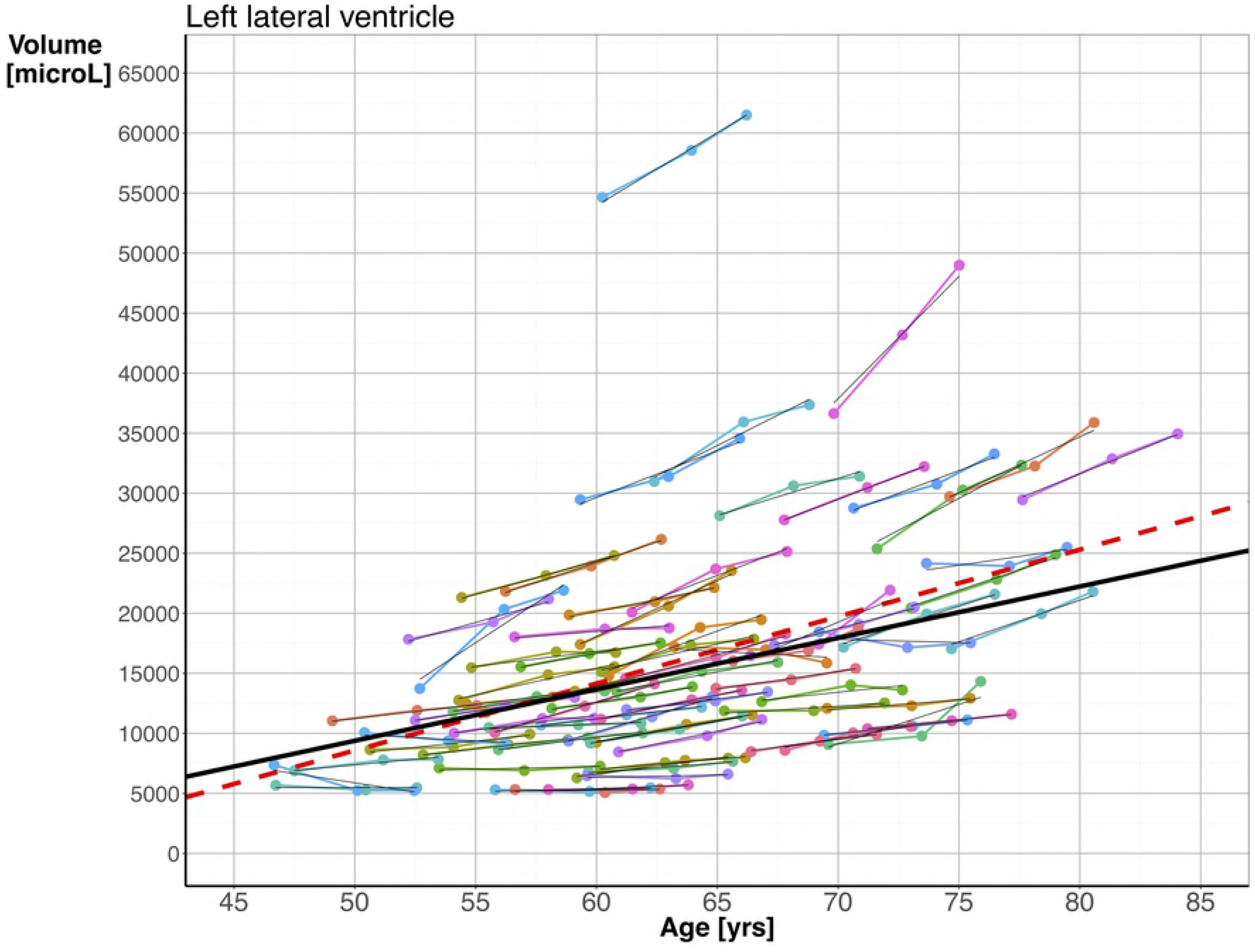
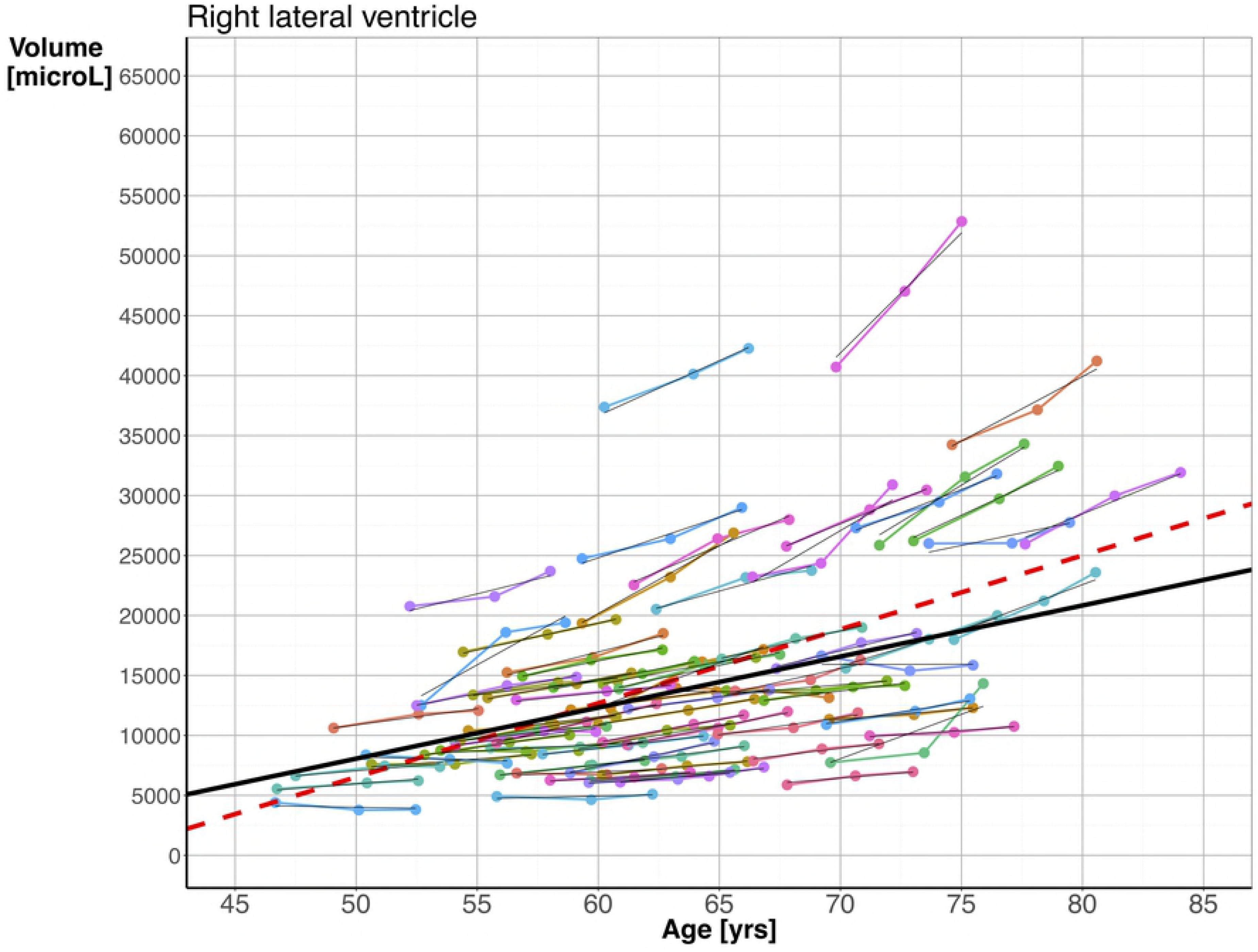
Subject-specific longitudinal lateral ventricle volumes versus age in left (a) and right (b) hemisphere shown as color-coded line plots across the three study waves. For left and right hemisphere the random effects, estimated from the linear mixed-effect model *Vol_ij_* = *β*_0_ + *β*_1_Age_*ij*_ + (*b*_0*i*_ + *b*_1*i*_Age_*ij*_) + *ϵ_ij_*, are depicted as thin line segments in black superimposed on the line plots. The thick regression line in black represents the estimated fixed effect, and the broken line represents ordinary linear least squares regression (OLS) line.

## 3 Results

### 3.1 Three wave changes in lateral ventricular volumes

A linear mixed-effect (LME) model was used to investigate the age-related evolution of the ventricular volumes. Figs 3a and 3b show the fixed and random effects lines calculated from the LV modeling for the left and right hemispheres, respectively. The fixed effects line shows expansion of LV volumes with increasing age, with an overall cohort change of 428.73 *μ*L/year for the LV on the left side, and 425.99 *μ*L/year for the right side. There was a trend towards a steeper slope in the oldest part of the sample, but the fixed effects line was less steep than ordinary linear least squares regression line (broken line), which did not take the dependencies between the measures from the three study-waves into account.

### 3.2 Explorative data analysis

The mean response inhibition performance (reaction-time) in the third study-wave was 56.99 s (sd = 14.5 s).

Figs 4 shows the distributions of age, the four volume measures, the response inhibition performance (RI), and their correlations. The influence of gender is illustrated by presenting the results separately for females and males. The LV volume measures for females were shifted to the lower end of the distribution, while the gender distributions were similar for age and RI. The univariate correlations between the left (*β*_1_L) and right (*β*_1_R) steepness measures (r = .94) and the two deviance measures VdevL and VdevR (*r* = .89) were strong. Statistically significant correlations between RI and the left (*r* = 48, p < .001) and the right (r = .53, *p* < .001) steepness measures of LV volumes were found for females only. The correlations between RI and the LV volume measures were non-significant.

**Fig. 4:**
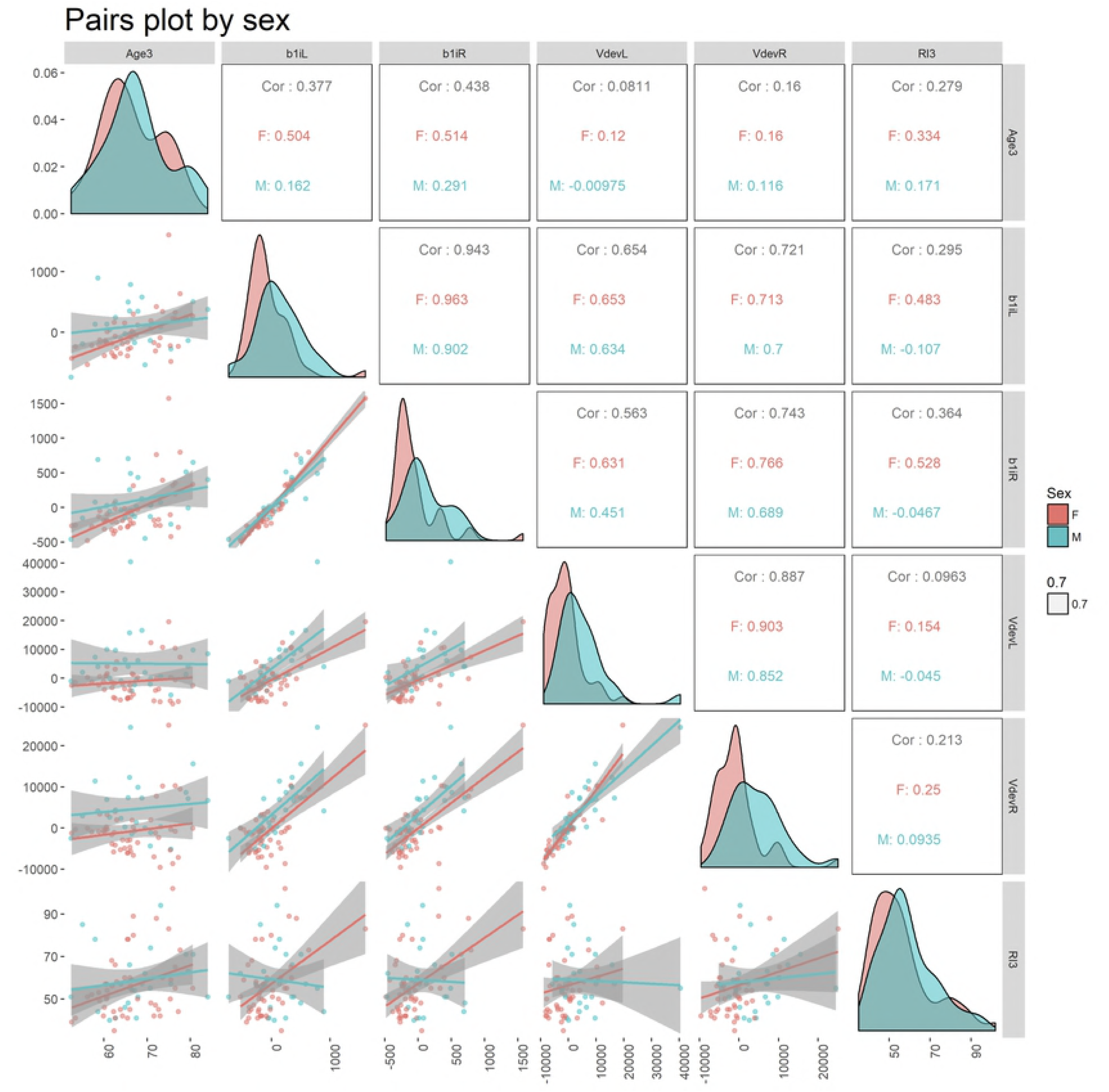
Pair-plot illustrating the distributions and correlations between age, the four LV volume trajectory measures and response-inhibition. The results are given separately for females (in red) and males (in green).

### 3.3 Predicting response-inhibition from trajectories of the LV volumes

The XGBoost analysis was used to investigate the relative importance of the four measures characterizing individual LV volume trajectories in predicting RI performance. Figs 5 shows that the most important feature was defined from the LV volume in the left hemisphere at inclusion. The second most important feature was selected from the right hemisphere, and represents the steepness of the trajectory from the first to the third study-wave. These two features were chosen for the final cross validation procedure.

**Fig. 5:**
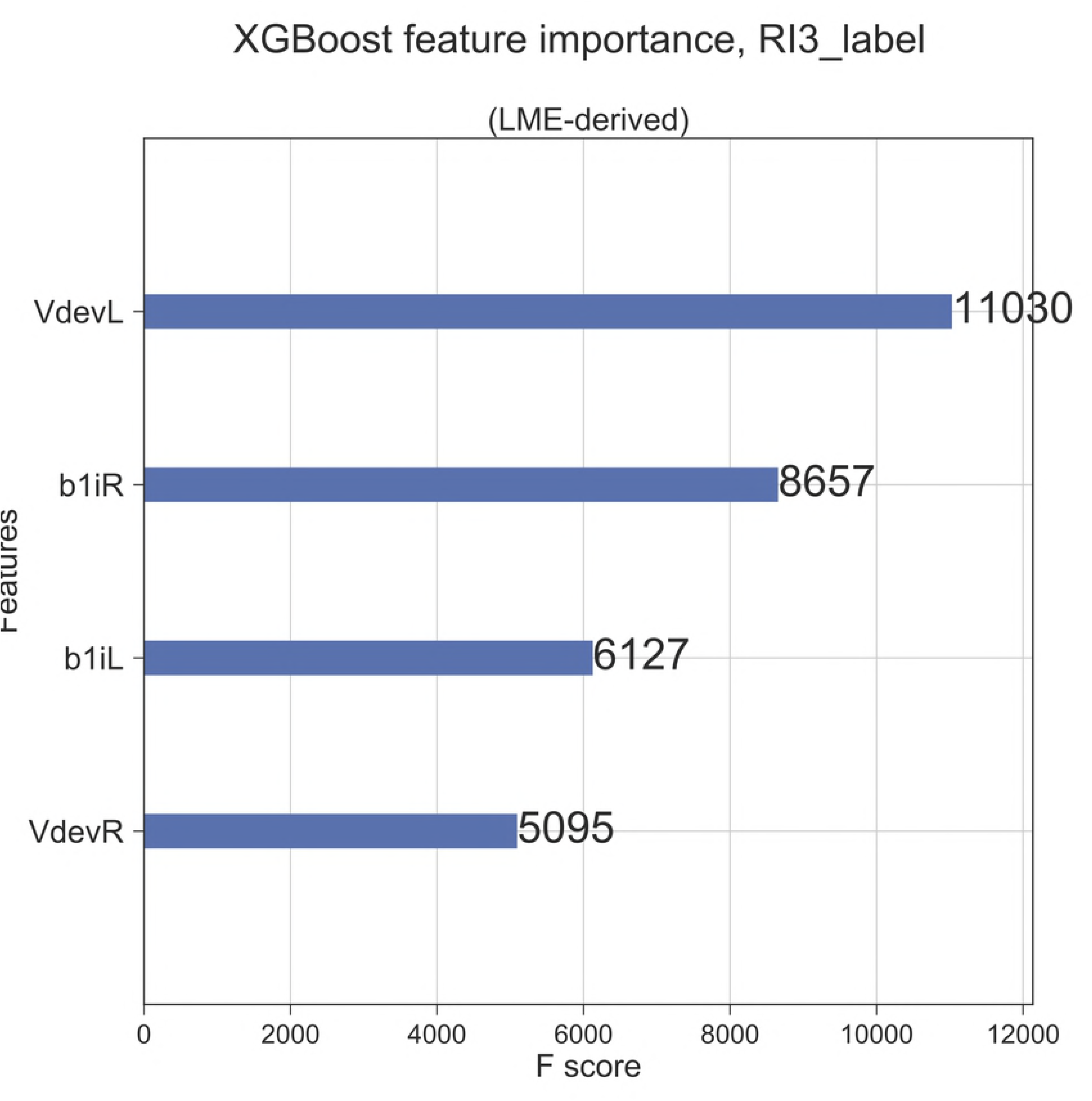
The Figure shows the result from the XGBoost analysis including the four selected measures of LV volume trajectories. VdevL/R: the deviance measure in the left/right hemisphere; b1iL/R = steepness of the trajectory in the left/right hemisphere.

A supplementary analysis was conducted to investigate the importance of gender when included together with the four LV volume measures and age in the XGBoost procedure. The results convincingly illustrated a minor importance of gender. Even age did not obtain stronger importance than the volume measures.

### 3.4 Prediction using a *k*-fold cross validation framework

Table 2 summarizes the classification performance using our cross-validation procedure including a linear multinominal logistic regression analysis, a nonlinear multilayer perceptron, and a voting classifier. According to the voting classifier, selecting the class label that represents the majority (mode) among the two other classifiers, the predictions of response inhibition category yielded an accuracy, precision and recall that were about .50. For this multinomial distributed sample divided into three classes with close to equal probabilities, this classification result is clearly above chance level. Taken together, the results support that a larger longitudinal expansion in the LV volumes is associated with poorer response inhibition in a given subject.

**Table 2:** Classification performance using the three different classifiers in stratified *k*-fold validation scheme (*k* = 10 splits) of the ventricular volumes evolution, estimated by the LME modeling. The features were standardized to zero mean and unit variance, and the performance measures represents the mean and standard deviation (SD) from the set of *k* folds. Each fold (test and train data splits) were kept the same for each classifier using the `scikit-learn` pipeline mechanism. MLR = multinominal logistic regression; MLP = multilayer perceptron (thre layers); Voting = the soft voting classifier predicting the class label based on the argmax of the sums of the predicted probabilities among the two other classifiers, MLR and MLP.

|  | Accuracy | Precision | Recall |
| --- | --- | --- | --- |
| MLR | 0.514 (0.179) | 0.483 (0.245) | 0.514 (0.179) |
| MLP | 0.417 (0.178) | 0.377 (0.198) | 0.417 (0.178) |
| Voting | 0.507 (0.179) | 0.499 (0.241) | 0.507 (0.179) |

## 4 Discussion

The present study showed an almost linear age-related expansion of the lateral ventricle volumes over a six years period. Four variables were selected from the linear mixed-effect analysis (LME) to characterize the individual trajectories in the sample: a random slope representing the steepness of change from the first to the third study-wave, and a measure of the LV volumes when included in the study, i.e., the distance between the observed volume and the volume expected from the age-related fixed cohort effect. For both variables, one was estimated for the right and one for the left hemisphere. The univariate exploratory data analysis showed that distributions of these four LV volume measures were characterized by gender differences, and that the response inhibition performance correlated strongly with age and the steepness measures in females. The multivariate XGBoost analysis did, however, show that age and gender were of minor importance compared to the four brain measures in predicting response inhibition performance. Two measures - the steepness measure from the right hemisphere and the deviance measure from the left hemisphere - were computed as the two most important features. When included in the 10-fold cross-validation procedure, they were shown to predict performance on the test of response inhibition clearly above chance level.

The trajectories of the lateral ventricle volumes depicted in Fig 2 confirmed results from previous studies showing that healthy aging is associated with expansion of the ventricular system [5,26]. The pair-wise correlation analysis showed that statistically significant results between the steepness measure and age were restricted to the female part of the sample. Gender differences have also been reported in previous studies on age-related LV volume changes [38,39], with age-related expansion described as slower in females than males. The deviance measures from the left and right hemisphere, selected as two features of the LV volume trajectories in our study, were lower in females than males across all ages. This reflects that the LV volumes were lower in females than males when included in the study. The effect of gender on the steepness measures did, however, show a different pattern: although females showed a lower steepness in the younger age groups, this trend shifted to a male dominance in the oldest part of the sample. This emphasize the importance of including a wide age range in studies of brain-behaviour relations, noting that the conclusions by Chung et al. [38] and Hasan et al. [39] included groups in their 40s and between 18 and 59 years, respectively.

The pair-wise correlation analysis did also reveal strong associations between response inhibition performance and the steepness measures of the LV volume trajectories. By this, the present results were similar to the results reported by McDonald et al. [13] and Aljondi et al. [14], assessing atrophy rates and corresponding changes in cognitive function in groups of older adults at two time-points. The conclusions from McDonalds et al. [13] were, however, based on results from a sample of patients with Mild Cognitive Impairment (MCI), and no significant correlations were found among any of the lobar atrophy rates and cognitive measures in the healthy controls. Aljondi et al. [14], using the same FreeSurfer longitudinal stream procedure as in the present study to estimate atrophy and a linear regression analysis to model brain-cognition changes, included a sample of healthy older women. In a ten-years follow-up study, they showed that atrophy rates were associated with increased rates of cognitive decline. Interestingly, our results from the explorative data analysis within the females subsample were similar to the results by Aljondi et al. [14].

Results from the explorative analysis thus suggest a gender specific brain-cognition relationship in older age. However, the correlation approach, including pairs of variables, are obviously not appropriate to detect the relative importance of demographic variables like gender. The machine-learning approach included in the present study rather demonstrated that gender were of minor importance when included together with the four measures of LV volume trajectories. Furthermore, although a pair-wise analysis showed weak correlations between response inhibition performance and the deviance measures, the importance of LV volumes when included in the study was clearly demonstrated by the machine-learning approach. When included in the cross-validation procedure together with the steepness measure of the LV volume change from the first to third study-wave, the prediction validity of response inhibition performance was clearly above chance level. The importance of level of performance have been widely discussed in studies of trajectories of change in cognitive function (e.g., [45]) and brain imaging (e.g., [1]), and separation of the effects of aging from differences at baseline is described as one of the many challenges when interpreting results from longitudinal studies. Our study showed the importance of considering this challenge by including information from baseline when selecting features characterizing trajectories of age-related changes, not at least in further studies inspired by a life-time perspective on brain health [46].

The statistical approaches selected in the present study have gained increased attention and support during the last two decades (see [47] for an overview), and we therefore believe that the present results may inspire further studies with a goal to obtain more accurate diagnostic and monitoring tools for brain health in older age. The continuous influence of such factors on the brain and cognitive function throughout the life-span, emphasize the importance of such studies also for detection of signs of Mild Cognitive Impairment (MCI), where a cognitive decline is described as more severe than expected from age and education level. The importance of detecting and treat patients with MCI is underscored by results showing that more than 50% of those defined as MCI are expected to progress to dementia within five years [48]. Currently, temporal lobe atrophy are commonly used as a diagnostic tool for neurodegenerative disorders.

The findings in this study give arguments for including information about changes in LV volumes, and by this supporting studies showing that such measures could even be more sensitive than a temporal lobe atrophy change rate to identify a neurodegenerative disorder (e.g., [49]).

There are several limitations in our study that need to be commented. The exclusive use of morphometric measures is one such weakness. Inclusion and fusion of multimodal MRI data (e.g., diffusen MRI and fMRI) is expected to provide a more comprehensive description of the brain patterns and a more accurate prediction of cognitive function. Inclusion of a larger set of features is, however, dependent on a much larger data set than in the present study, and gives a strong argument for sharing data in future studies [50]. Inclusion of performance on only one cognitive test may be considered as another weakness. Previous studies on brain-behaviour relations in the aging brain have mainly focused on episodic memory function [12]. We will argue that a measure of response inhibition, known to be affected as part of normal cognitive aging [23,24], should not be underrated as an important outcome measure. The so-called ”ability impurity” [25] of this and other measures of executive function can rather be evaluated as a strength, in that the performance is likely also dependent on abilities that are commonly defined within other cognitive domains (e.g., processing speed and memory function). To sort this out, future studies should include both more “pure” measures and more general composite measures. Finally, the selection of a general brain measure like LV volumes can be criticized. Most studies have included more specific brain regions. However, the association between a given cognitive domain and a regional brain area is not clear-cut, which can be illustrated by studies associating structural measures of hippocampal volume with episodic memory function [51], executive function [52] as well as processing speed [53]. There are obviously many factors influencing brain measures like LV volumes as well as different aspects of cognitive function trough the life-span, and even interactions between brain-behaviour measures in more general [54]. Although we did find the clear pattern of our LV volume measures intriguing, future studies should follow the advice from Steffener et al. [54] to utilize multiple neuroimaging modalities within a conceptual model of cognitive aging when predicting complex function defined within the concept of executive function and cognitive control.

Taken together, the results in the present study confirmed the importance of brain changes for cognitive function in older age. Individuals differ both regarding brain changes and the associated pace and level of cognitive changes in older age [1], leaving us with a task to search for ways to preserve our brain. A study by Habeck et al. [55] used a computational model to quantify the brain maintenance and cognitive reserve for single subjects by including chronological age, neuropsychological performance and structural brain measures as features. In a follow-up study they identified cognitive reserve networks that was task irrelevant, supporting that life experiences may preserve the individual against age- or disease related changes [56] We believe that inclusion of such features into a predictive model similar to the one used in the present study would give further information about factors of importance to brain health and successful cognitive aging in older age [12].

## 4.1 Acknowledgement

The first study-wave was financially supported by the Research Council of Norway (grant 154313/V50), the second and third study-wave from the Western Norway Regional Health Authority (grants 911397 and 911687). We also acknowledge the Western Norway Regional Health Authority for funding the Freesurfer Longitudinal Stream (grant 911995), and the Bergen Research Foundation for support through the project ”Computational medical imaging and machine learning - methods, infrastructure and applications”. Finally, we thank colleagues and participants in the longitudinal study on cognitive aging.

